# Inferred Model of the Prefrontal Cortex Activity Unveils Cell Assemblies and Memory Replay

**DOI:** 10.1101/028316

**Authors:** Gaia Tavoni, Ulisse Ferrari, Francesco P. Battaglia, Simona Cocco, Rémi Monasson

## Abstract

Cell assemblies are thought to be the units of information representation in the brain, yet their detection from experimental data is arduous. Here, we propose to infer effective coupling networks and model distributions for the activity of simultaneously recorded neurons in prefrontal cortex, during the performance of a decision-making task, and during preceding and following sleep epochs. Our approach, inspired from statistical physics, allows us to define putative cell assemblies as the groups of co-activated neurons in the models of the three recorded epochs. It reveals the existence of task-related changes of the effective couplings between the sleep epochs. The assemblies which strongly coactivate during the task epoch are found to replay during subsequent sleep, in correspondence to the changes of the inferred network. Across sessions, a variety of different network scenarios is observed, providing insight in cell assembly formation and replay.

**Author Summary:** Memories are thought to be represented in the brain through groups of coactivating neurons, the so-called cell assemblies. We propose an approach to identify cell assemblies from multi-electrode recordings of neural activity *in vivo,* and apply it to the prefrontal cortex activity of a behaving rat. Our statistical physics inspired approach consists in inferring effective interactions between the recorded cells, which reproduce the correlations in their spiking activities. The analysis of the effective interaction networks and of the model distributions allows us to identify cell assemblies, which strongly co-activate when the rat is learning, and also during subsequent sleep. Our approach is thus capable of providing detailed insights in cell-assembly formation and replay, crucial for memory consolidation.

## Introduction

Cell assemblies [1, 2, 3], closely connected, synchronously activating groups of cells have been posited as the main constituents of memory and information representations. The activation and reactivation (‘replay’) of cell assemblies is thought to be critical for consolidation and re-elaboration of memories, working memory and decision making [4, 5, 6]. The precise characterization of cell assemblies from experimental data remains, however, very difficult. Brute force and exhaustive search for groups of neurons with strongly correlated firings is impossible due to the combinatorial number of possibilities. Current available methods for cell assembly detection and replay estimation often rely on the identification and on the matching of templates [7, 8, 9]. In the hippocampus, for instance, such templates are provided by the temporal sequence of firing events of place cells during the awake phase. The correlational structure of data can also be used to approximate templates from principal component analysis [10, 11], or to search for clusters of neurons with related firing patterns [12, 13].

In this work we propose an alternative approach for identifying cell assemblies. We first infer a model for the distribution of the neural activity from the recorded spiking activity, based on an estimate of the effective coupling network between the neurons (Fig. 1). While the anatomical synaptic structure is usually unknown [14], the effective couplings may be inferred by methods borrowing from statistical inference and statistical physics [15], mapping the observed spiking data onto abstract network models, *e.g.* generalized linear models [2, 16, 17], integrate-and-fire models [18, 19], Boltzmann machines [20] and Ising models of binary neurons [21, 22, 23, 24, 25]. The effective-coupling-based model allows us to characterize the firing probability of any neuron conditional to the activity of the other cells in the population. We may then search for *self-sustaining activity patterns,* in which each neuron ‘reads’ the activities of the other recorded cells, and, in turn, participates as an input to those cells in a coherent way. This concept of self-sustaining pattern is closely related to Hebb’s classical definition of a cell assembly as *”a diffuse structure … capable of acting briefly as a closed system”* [1]. Self-sustaining patterns are encoded in the effective-coupling network, arise spontaneously whenever favorable network or cellular excitability are met, and may be detected by a downstream ‘reader’ neuron [3]. A computationally-efficient way to search through the combinatorial number of putative self-sustaining patterns consists in applying a driving input favoring high-activity configurations, mimicking transient increases in network excitability, as may be induced by e.g. slow oscillations of sleep. As the drive gets stronger self-sustaining activity patterns with more and more active neurons are revealed (Fig. 1). Those patterns contain groups of strongly coactivating neurons, which define our cell assemblies.

**Figure 1:**
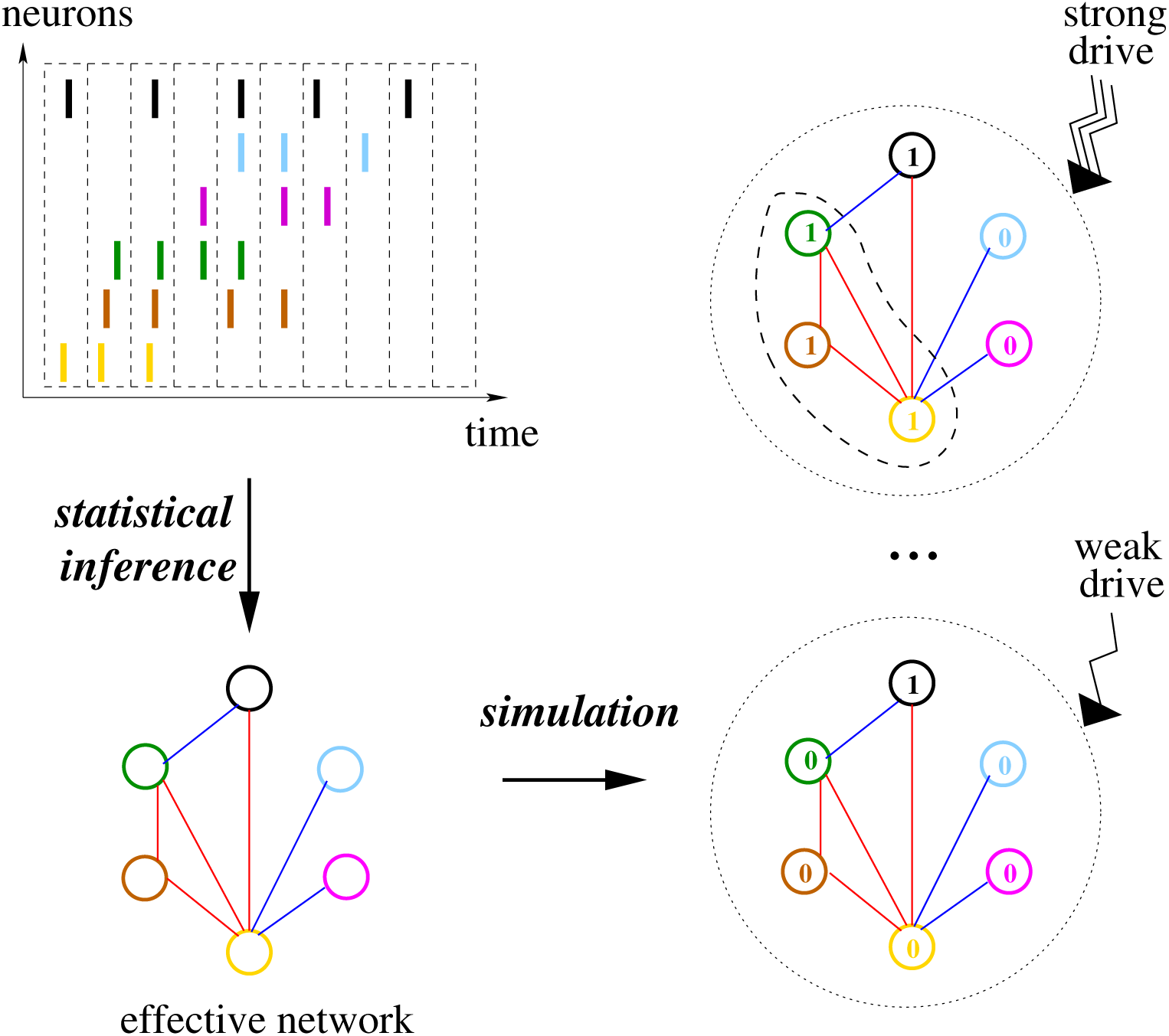
Model for the neural activity: definition and simulation. Spiking times are binned into time bins of width Δ*t*; each neuron *i* is assigned the variable *σ_i_* = 1 or 0, if it is active in the time bin or not (**top, left**). A model of the neural activity distribution (*P* in Eq. [1]) is inferred to reproduce the 1- and 2-cell firing frequencies of this binned data (**bottom, left**); Red and blue links correspond, respectively, to positive and negative effective couplings *J_ij_* in the inferred network. The model distribution is then simulated with the addition of an increasing drive, which favors configurations with more and more active neurons; Values of the activity variables *σ_i_* in the most likely configuration given the drive are shown (**right**). As the drive increases a group of neurons (comprised in the dashed contour) may abruptly coactivate, defining a cell assembly.

Here, we infer the effective couplings of Ising models [26] that best reproduce the distribution of activity of prefrontal neurons during performance of a decision-making task [10, 27] and during the preceeding and following sleep epochs. Comparison of the sleep interaction networks reveals the presence of task-related effective potentiation or depression of the effective couplings. We then simulate the models of each epoch under an external drive, thereby identifying self-sustaining configurations of neural activity permissible at different levels of neural excitability, and putative cell assemblies. The cell assemblies supporting the effectively potentiated couplings are shown to strongly coactivate in the behavioral epoch but not in the preceding sleep epoch, and to be replayed in the subsequent sleep epoch. A wide-scale study of about 100 experimental sessions shows a variety of possible scenarios for the cell assemblies across the epochs, which allows us to formulate empirical rules for their formation and replay.

## Results

### Model for the neural activity distribution and effective coupling network

We briefly present the approach to model the distribution of activity of the *N* recorded neurons (see Methods for more details). The spiking times are binned within small time bins of duration Δ*t* = 10 ms; the activity configuration (*σ*_1_,σ_2_*, …,σ_n_*) are snapshots of the neural activity, where *σ_i_* takes values one or zero depending on whether the *i*-th neuron is, respectively, active or inactive in the time bin. We model the probability distribution of activity configurations as

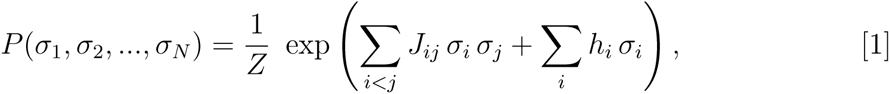

where *Z* ensures normalization of the distribution. The 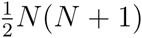 parameters *h_i_* and *J_ij_* are fitted to reproduce the *N* individual spiking frequencies *f_i_* and the 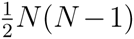 pairwise spiking frequencies *f_ij_* (within a time-bin Δ*t*) estimated from the recording data, see Methods. *P* in Eq. [1], called Ising model in statistical physics, is the least constrained (with maximum entropy), default probability distribution reproducing this low-order spiking statistics [21].

Parameters *J_ij_* define the effective pairwise couplings between the cells (Fig. 1): *J_ij_* different from zero expresses the presence of a conditional dependence between neurons *i* and *j*, not mediated by other neurons in the recorded population. The conditional average activity of neuron *i* given the other neuron activities {*σ_j_*}, with *j ≠ i*, reads

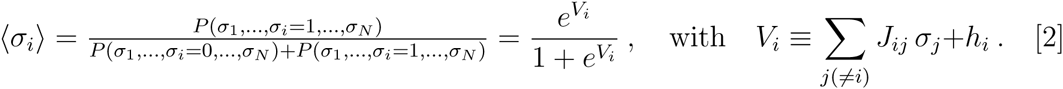

It is a logistic function of its total input, *V_i_*, equal to the sum of the other neuron activities *σ_j_* weighted by the couplings *J_ij_*, and of the local input *h_i_*.

We have re-analyzed recordings of the activity of tens of neurons in the pre-frontal cortex of five behaving rats [10]. Each recording session is divided in three ≃ 30-minute epochs: a Task epoch in which the rat had to learn a rule (go left, right, where the light is on, or off, in a Y-shaped maze), which was changed as soon as the rat had learned it, and two Sleep epochs, one before (Sleep Pre) and one after (Sleep Post) the Task epoch. We have inferred with the adaptive cluster expansion of [26] the parameters *h_i_* and *J_ij_* for the three epochs of the 97 recorded sessions, together with their statistical error bars, Δ*h_i_* and Δ*J_ij_* (Methods). The inferred model distribution *P* reproduces the single-neuron and pairwise spiking probabilities in a time bin with great accuracy (Fig. 2a). In addition, it also predicts the value of higher-order moments such as triplet firing probabilities, and the probability of multiple neuron firing in a time bin, in excellent agreement with the data (Fig. 2b). Our model approach successfully complies with standard criteria for statistical inference, such as cross-validation (Methods and Fig. 2c). In addition, the structure of the inferred interaction network is found to be largely sparse, with an average of about 60% of zero couplings across epochs and sessions, while about 40% of pairwise correlations are compatible with zero within one standard deviation. The Ising model therefore offers an accurate and compressed representation for the empirical distribution of activity snapshots, over an extended period of time.

**Figure 2:**
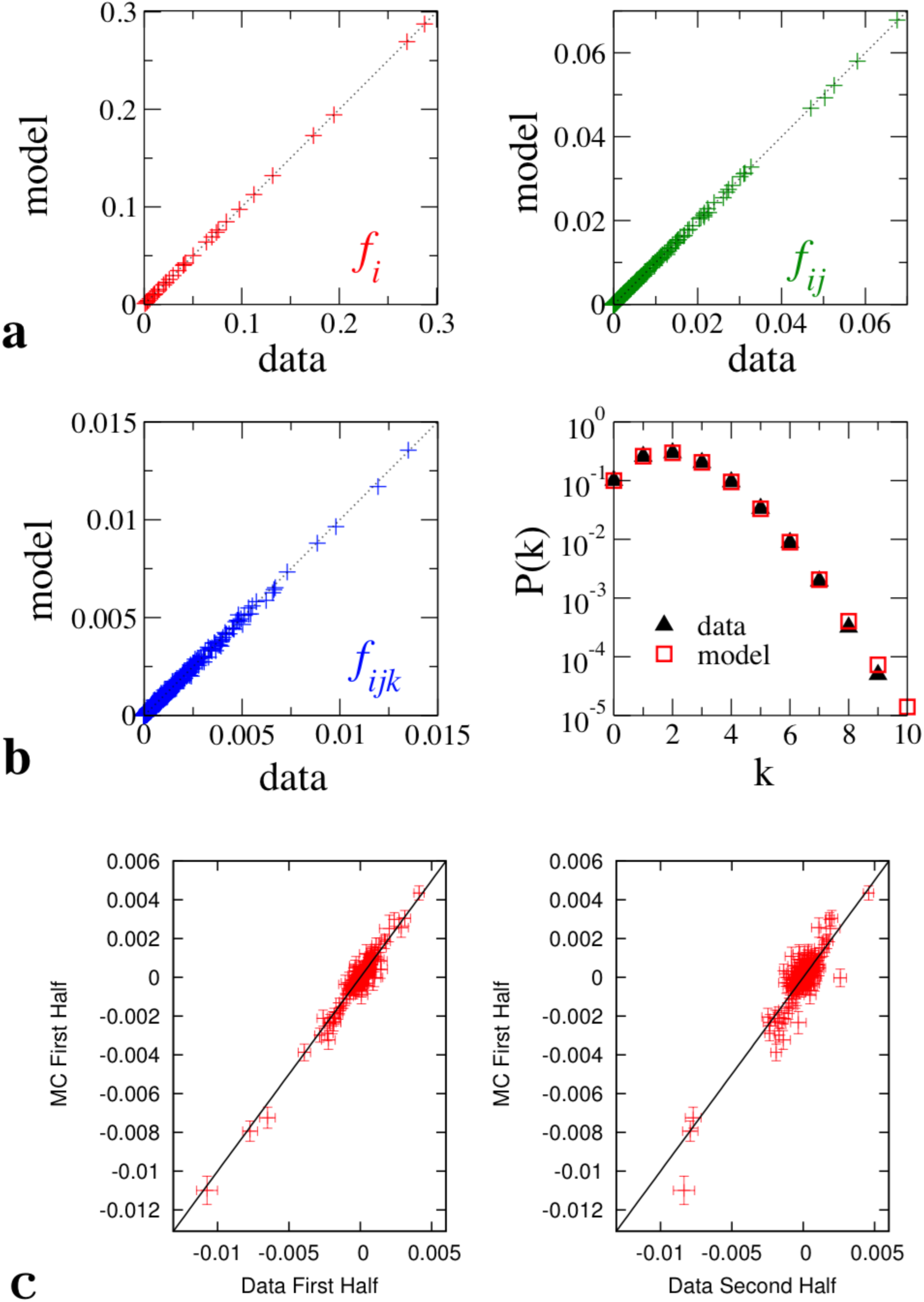
Quality and validation of the inferred model. **a**. Reproduction of the low-order statistics of the spiking data. Scatter plots of the single-neuron (*f_i_*, left panel) and of the pairwise (*f_ij_*, right panel) frequencies. Values of the frequencies computed from the spiking data are shown along the *x*-axis, while their counterparts computed from the inferred model distribution P, Eqn. [1], are shown along the *y*-axis. **b**. Predictions for higher-order statistics. Left panel: scatter plot of the triplet frequencies *f_ijk_* computed from the data (*x*-axis) and from the inferred model distribution (*y*-axis). Right panel: probability *p*(*k*) that *k* neurons are active in a time-bin (of duration Δ*t* = 10 ms), computed from the data and from the model distribution. The agreement is excellent for *k* such that *p*(*k*) times the number of time-bins is larger than or equal to one, that is, provided the recording time is sufficient to sample those rare configurations of multiple neuron firing. **c**. Cross-validation of the model distribution *P*, inferred from the spikes emitted in the first half of the recording of the Task epoch in Session A (Methods). The correlations *c_ij_ = f_ij_ – f_i_f_j_* are shown along the *x*-axis (left panel: first half of the recording, right panel: second half), and compared to the values computed from the model distribution (*y*-axis). In both panels the points lie close to the diagonal line, within one or two error bars corresponding to the statistical standard deviation due to the finite sampling. Moreover the offsets from the diagonal are of the same order of magnitude in the two panels, confirming the absence of overfitting in our method.

### Comparisons of Sleep epoch coupling networks reflect Task-related changes

We describe in detail the findings for one typical experimental session, called A, with *N* = 37 neurons, and we will present results on the remaining sessions in what follows. We first compare the networks inferred from the three recorded epochs of session A. The scatter plot of the inferred couplings is shown in Fig. 3, giving information about how couplings change between the epochs. Most couplings do not vary much between Sleep Pre, Task, and Sleep Post, while some are strong in the Sleep epochs only. Interestingly, some couplings are weak or negative in Sleep Pre, and become stronger and positive in Task and in Sleep Post (Fig. 3). In session A those effectively potentiated couplings are mostly supported by a group of five neurons (1-9-20-21-26 in Fig. 4, top panels), which are strongly and positively interconnected in Task and in Sleep Post, but not in Sleep Pre.

**Figure 3:**
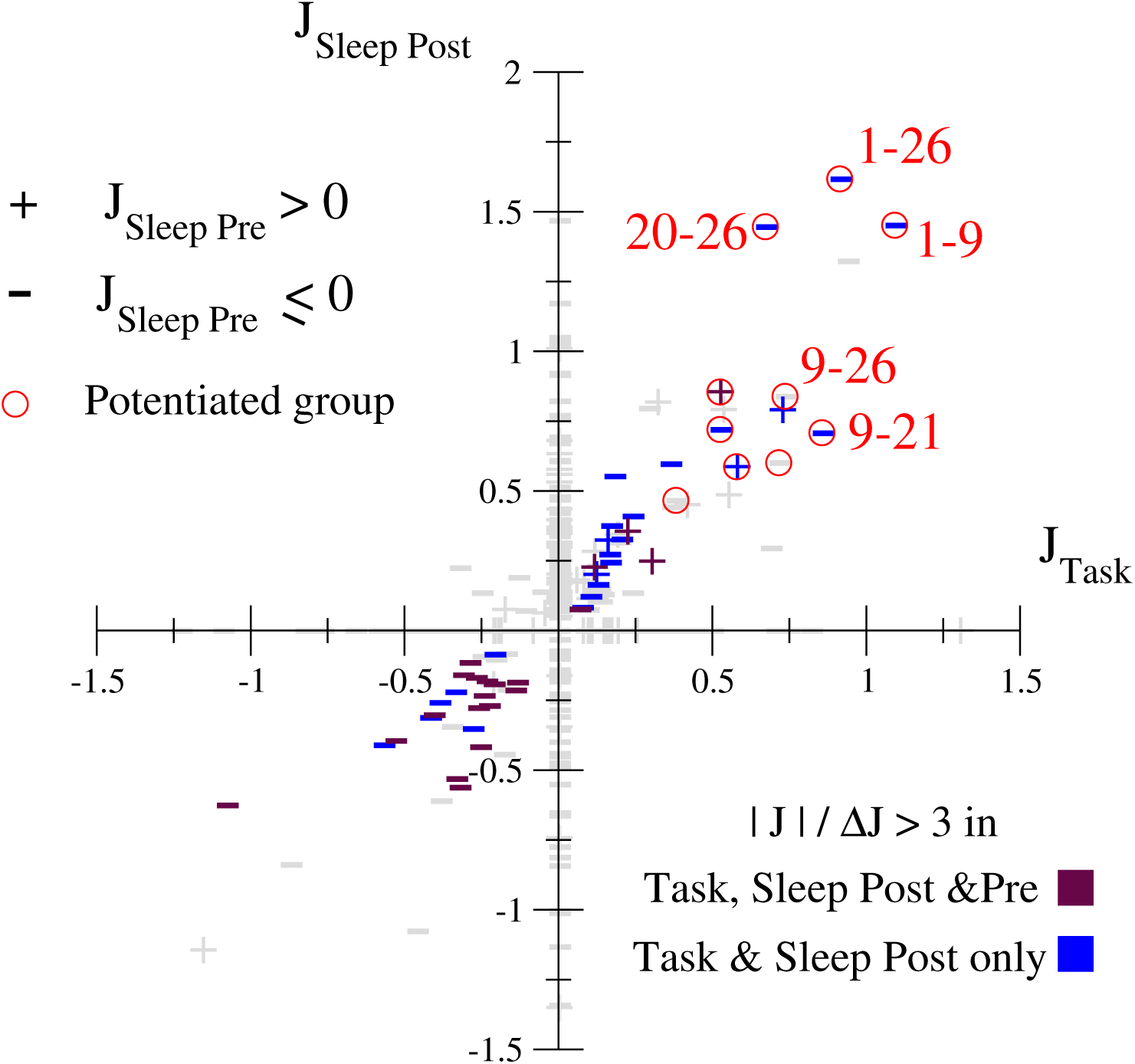
Comparison of the inferred couplings in the different epochs of Session A. Scatter plot of the Ising couplings inferred for the Task vs. the Sleep Post epochs in session A. Positive and negative (or null) couplings in the Sleep Pre epoch are shown with, respectively, + and − symbols. A group of five neurons (1-9-20-21-26) supports most of the potentiated couplings, shown by red circles; five of the pairs are shown with the corresponding neuron numbers.

**Figure 4:**
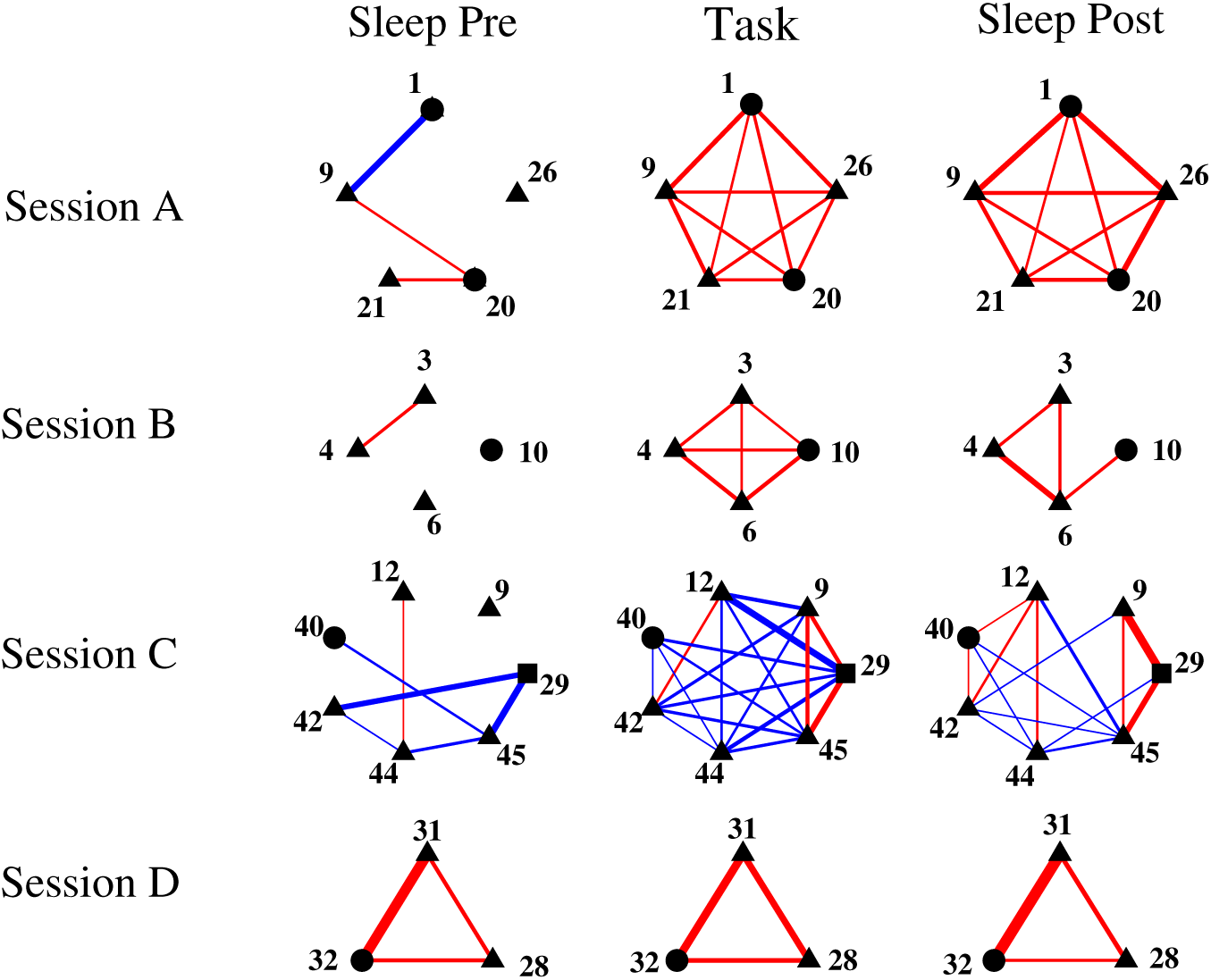
Relevant subnetworks of couplings in sessions A, B, C, D. Relevant subnetworks of couplings in the three epochs of sessions A to D (red: *J* > 0, blue: *J* < 0; Line thickness is proportional to |*J*|). Pyramidal cells are shown with triangles, undetermined cells with circles, and interneurons with squares. In Session A to C, we show the subnetworks of couplings with most changes between Sleep Pre and Post; the subnetworks in Sleep Post show large similarities with the ones in Task. In Session D, where no significative change in the inferred couplings is observed between Sleep Pre and Post, we show the most interconnected subgroup of (3) neurons. *Session A:* The potentiated group is composed of neurons 1-9-20-21-26 identified in Fig. 3. *Session B:* The potentiated group is composed of neurons 3-4-6-10. *Session C:* neurons in a potentiated subgroup (9-29-45) have inhibitory connections with the group (12-40-42-44). *Session D:* One 3-cell, largely connected group (28-31-32) is conserved across all three epochs.

Similar scatter plots can be drawn and studied for all available sessions. Figures corresponding to six representative sessions, labelled B to G, can be found in Supporting Information-II. Session B, with *N* =10 recorded cells only, displays a behavior similar to session A (Supporting Information, Fig. S10), with an effectively potentiated group of four cells (Fig 4). Session C exhibits a more complex network reconfiguration between the Sleep Pre and Post epochs, with the appearance of new positive (as in sessions A and B) and new negative couplings (Fig. S13). As shown in Fig. 4 the effectively potentiated group is made of three neurons (9-29-45), with couplings vanishing in Sleep Pre but large and positive in Task and Sleep Post. In addition, many couplings between this group and another group of four neurons (12-40-42-44) are strongly depressed, decreasing from zero values in Sleep Pre to negative values in Task and Sleep Post. In Session D no signature of task-related change in the couplings is found (Fig. S18); the three largest positive couplings define the network shown in Fig. 4, which is conserved across the three epochs. Session E includes two strongly potentiated couplings between two unrelated pairs of neurons (Fig. S21). Sessions F and G show very similar behaviors to, respectively, A and B, see Figs. S23 & S28.

The examples above show that Task-related changes of the couplings between the Sleep epochs greatly vary across the sessions. To quantify this effect in a way allowing us to scan efficiently the 97 sessions we introduce the following session-wide estimator:

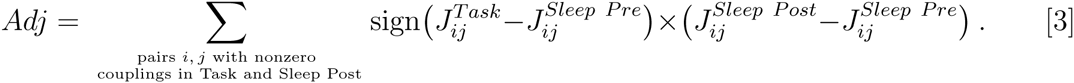

*Adj* is measure of the task-related *adjustment* of the inferred couplings between the Sleep Pre and Post epochs. The presence of the sign function allows us to sum constructively contributions corresponding to effective potentiation (as in sessions A, B, C) and depression (as in session C) of the couplings. The summation is restricted to pairs *i, j* of neurons, whose couplings are significantly different from zero in Task and Sleep Post. In practice we require that |*J_ij_*|/Δ*J_ij_* > 3, where Δ*J_ij_* is the error bar on the inferred coupling *J_ij_*, though the value of *Adj* varies little upon relaxing the criterion to |*J_ij_*|*/*Δ*J_ij_ >* 2.

Figure 5, left panel, shows the values of *Adj* vs. the numbers *N* of recorded cells for the 97 sessions. Some sessions (including, but not restricted to, A, B, C, E, F, G) have large and positive *Adj*, more than one standard deviation above a null model (where the correspondence between pairs of neurons across the epochs is removed by reshuffling the neuron indices, see Supporting Information-I.D) average. The outcome of a control calculation, where, for each session, we exchange the Sleep Pre and Sleep Post couplings in Eq. [3] is shown in Fig. 5, right panel. As expected no large–*Adj* session is found. This simple control provides a clear evidence for the fact that *Adj* captures experience-related changes in the couplings.

**Figure 5:**
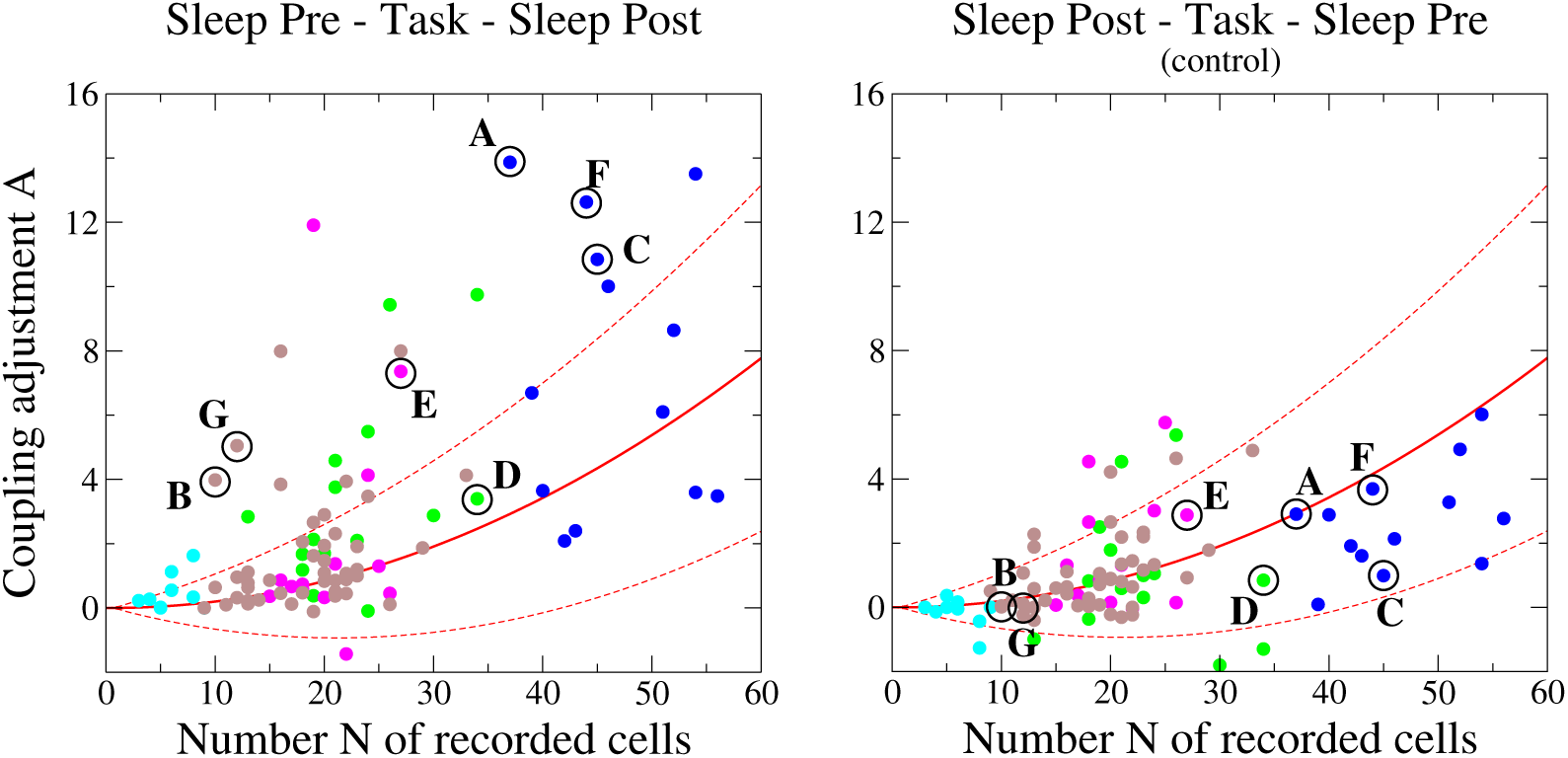
Coupling Adjustment across all 97 recorded sessions. Coupling Adjustment *Adj*, see Eqn. [3], is shown for the 97 sessions in the left panel. The right panel shows a control calculation, where we have exchanged the Sleep Pre and Post inferred couplings. Red lines show the predictions of the null model (average: full lines, ±1 standard deviation: dashed lines), see SI, Section I.D. Colors identify the five recorded rats; circles locate sessions A to G.

### Simulations of the inferred distribution of activity identify putative cell assemblies

We now simulate the model distribution *P* derived above to identify groups of neurons, or ‘putative cell assemblies’ most likely to co-activate. We call *self-sustaining* a configuration of activity (*σ*_1_, *σ*_2_, …, *σ_N_*) such that for each cell *i*, the total input *V_i_* in Eq. [2] is positive and the neuron is active (*σ_i_* = 1), or *V_i_* is negative and the neuron is silent (*σ_i_* = 0). In other words, in a self-sustaining configuration, each neuron activity is consistent with the inputs coming from the other neurons (and from *h_i_*). It is easy to show that self-sustaining configurations are local maxima of the distribution *P* (Methods), and are therefore prototypical patterns of the neural activity.

As the effective couplings *J_ij_* and local inputs *h_i_* reflect the neural activity distribution over the entire epoch, neural activity is quite sparse. A simulation of the model distribution with the parameters as inferred will not generate any self-sustaining state, but the all-silent neuron configuration, *σi* = 0 for all *i* = 1,…,*N*. In neural activity, cell assemblies are not sustained for a long period, but happen transiently during bouts of high neural excitability. To simulate that, we introduce an extra parameter *H* which increases the likelihood for neurons to activate (in an uniform way [28]) and allows us to reproduce assembly-generating transients. In practice, for each epoch of the recorded sessions, we identify the self-sustaining configurations upon changing the total inputs *V_i_* into *V_i_* + *H* in Eq. [2] for all neurons *i* (Methods). Figure 6 shows the number of active neurons in the self-sustaining configurations as a function of *H* for the three epochs of Session A. As *H* increases from zero, neurons start to activate one after the other, in decreasing order of their local inputs *h_i_*. As more and more neurons *i* get activated contributions to the total inputs of the other neurons *j* build up, facilitating (*J_ij_* > 0) or hindering (*J_ij_* < 0) their activations.

**Figure 6:**
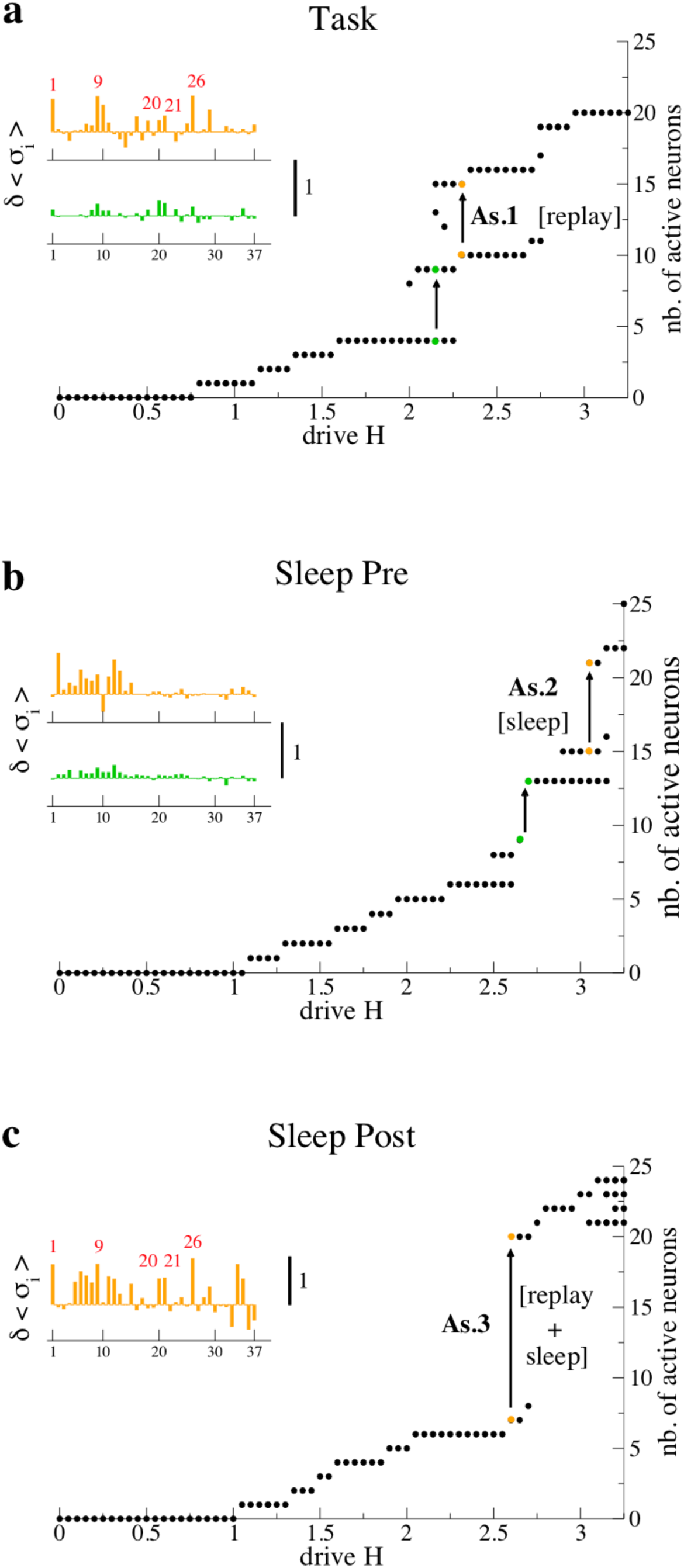
Identification of groups of coactivated neurons in Session A. Number of active neurons in the self-sustaining configurations of the Ising model as a function of the drive *H* for the Task (**a**), Sleep Pre (**b**), Sleep Post (**c**) epochs. Distinct self-sustaining configurations may coexist (at a given *H*) and are indicated by the same colored dots, and defines jumps indicated by arrows. The changes in the conditional averages 〈*σ*_i_〉 corresponding to the self-sustaining configurations shown by the colored dots are given in insets. While the first two jumps in Sleep Pre and Task correspond to small changes in 〈*σ*_i_〉, the second two jumps in those epochs and the jump in Sleep Post include a group of neurons with substantial changes in 〈*σ*_i_〉, which define our cell assemblies. As.1 includes neurons 1-9-10-16-18-20-21-26-29, As.2 includes 2-6-7-8-9-11-12-13, and As.3 includes 1-5-6-7-8-9-11-12-15-20-21-26-29-34-35. Two groups of neurons, common to the cell-assemblies of the different epochs, are the replay (1-9-20-21-26) and the sleep (6-7-11-12) groups.

Discontinuous ‘jumps’ in the number of active neurons are found at special values of *H*, and are indicated with arrows in Fig. 6. A jump signals the coexistence of two self-sustaining configurations, with, respectively, a low and a high numbers of active neurons. We take such co-existence of a high and a low-activity configurations as the hallmark of a cell assembly, as it highlights how transiting to a new self-sustaining state requires the concomitant activation of a group of neurons. We show in the insets of Fig. 6 the variations *δ*〈*σ_i_*〉 in the conditional average activities, 〈*σ_i_*〉 in Eq. [2], between the ‘low’ and ‘high’ self-sustaining configurations. As the local inputs *h_i_* and the drive *H* are constant across the jump large variations *δ*〈*σ_i_*〉 (in absolute value) may come only from the collective activation of a group of neurons *i* (Fig. 1 & 4), which we call putative cell-assembly (As), see details in Methods and Supporting Information-II.A. In either Task or Sleep Pre only one out of the two jumps defines a putative cell assembly, while the other jump does not show strong variations *δ*〈*σ_i_*〉, see Fig. 6a & 6b and Insets. In Sleep Post the single jump defines a large putative cell assembly (Fig. 6c).

Comparing the cell assemblies of session A across the different epochs we find common subgroups of neurons (Fig. 6 and caption). The ‘Sleep’ group (neurons 6-7-11-12) is shared by As.2 in Sleep Pre and the large assembly As.3 in Sleep Post (Figs. 6b&c). The subnetworks of couplings, inferred in the three epochs of session A, between the neurons in the Sleep and Replay groups are shown in Fig. 7. Neurons in the Sleep group are the only ones to be interconnected by large and positive couplings in the Sleep Pre epoch, and the same statement holds for the cells in the Replay group in the Task epoch. In the Sleep Post epoch, however, the two groups becomes largely interconnected. The merging of the Sleep and Task cell assemblies in a large, interacting group of cells in Sleep Post, corresponds to the large coactivation jump in Fig. 6c, is a general finding for all the sessions with a large coupling Adjustment, see description of scenarios below.

**Figure 7:**
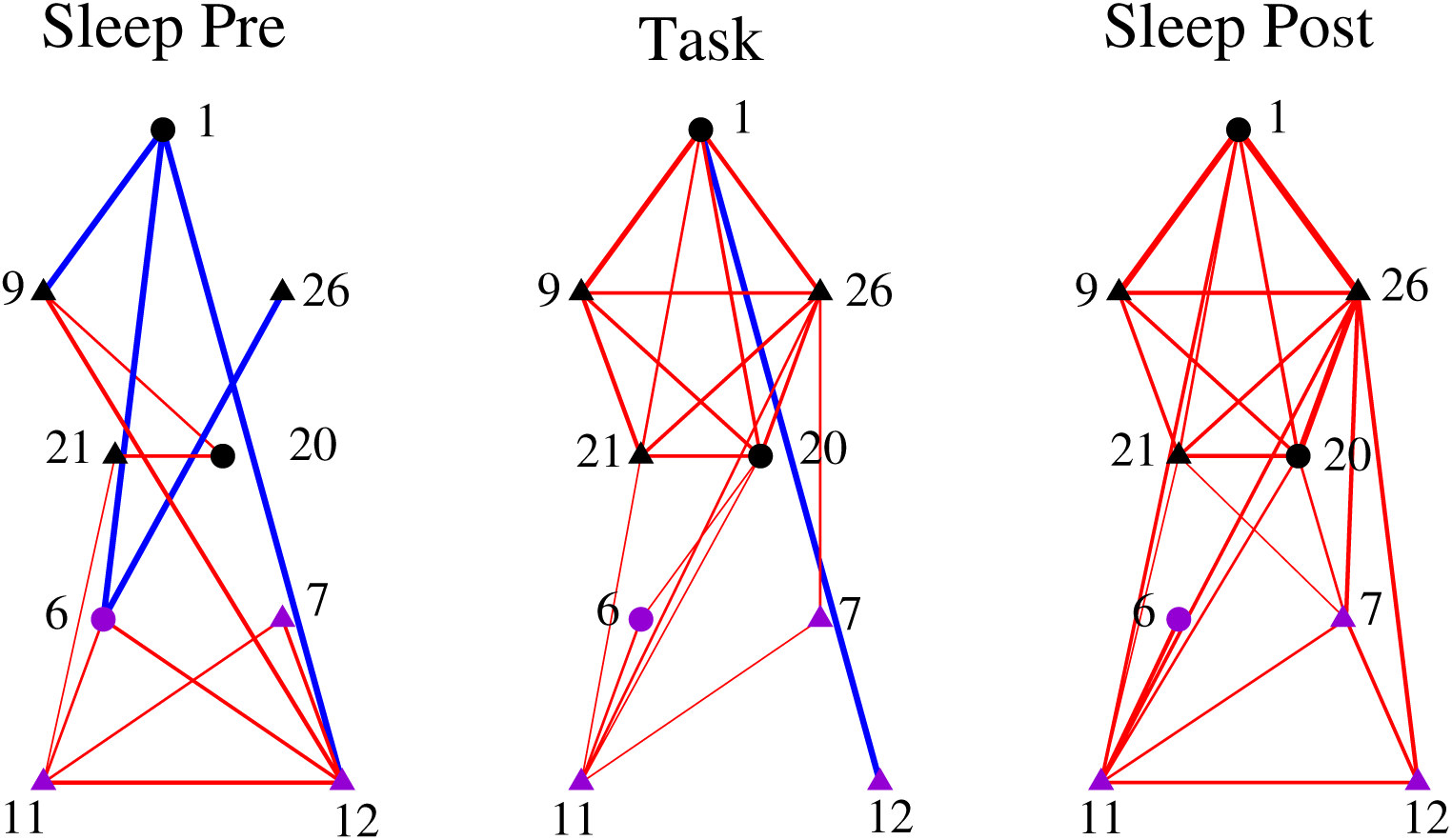
Association of the Replay- and Sleep-group subnetworks in the Sleep Post epoch of session A. Subnetwork of strong couplings within the Replay (top five neurons 1-9-20-21-26, black symbols) and Sleep (bottom four neurons 6-7-11-12, violet symbols) groups identified in Fig. 6. Red: *J* > 0, Blue: *J* < 0. Line width is proportional to the coupling intensity. Pyramidal cells are shown with triangles.

### Identified Task-related groups of coactivating neurons replay during subsequent Sleep

We now show that the putative assemblies found in the model simulations correspond to real coactivations of the attached neurons in the spiking data. To this aim we define the co-activation ratio (CoA) of a group *G* of neurons over the time scale *τ* through

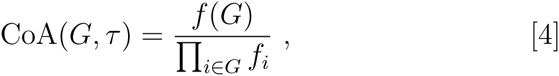

where *f*(*G*) is the probability that all the neurons in the group are active within the time scale *τ*, and the denominator is the product of the individual spiking probabilities. For a group of independent cells the CoA is on average equal to unity. As a relevant coactivation event contributing to *f*(*G*) should correspond to a sequence of ’readings’ of spikes, triggering in turn the next spike, we expect *τ* to be not larger than *n ×* Δ*t*, where *n* is the number of neurons in *G* and Δ*t* = 10 ms is the time-bin duration used for the inference.

We have computed the CoA of the cell assembly As.1 from the spiking data, with the results shown in Fig. 8a. As.1 is found to strongly coactivate in Task and in Sleep Post (on much longer time scales), but only during Slow-Wave-Sleep periods (SWS), in which hippocampal sharp waves are known to be important for memory consolidation. In Sleep Pre As.1 does not coactivate, which is compatible with the independent-cell hypothesis due to the low firing frequencies (Methods).

**Figure 8:**
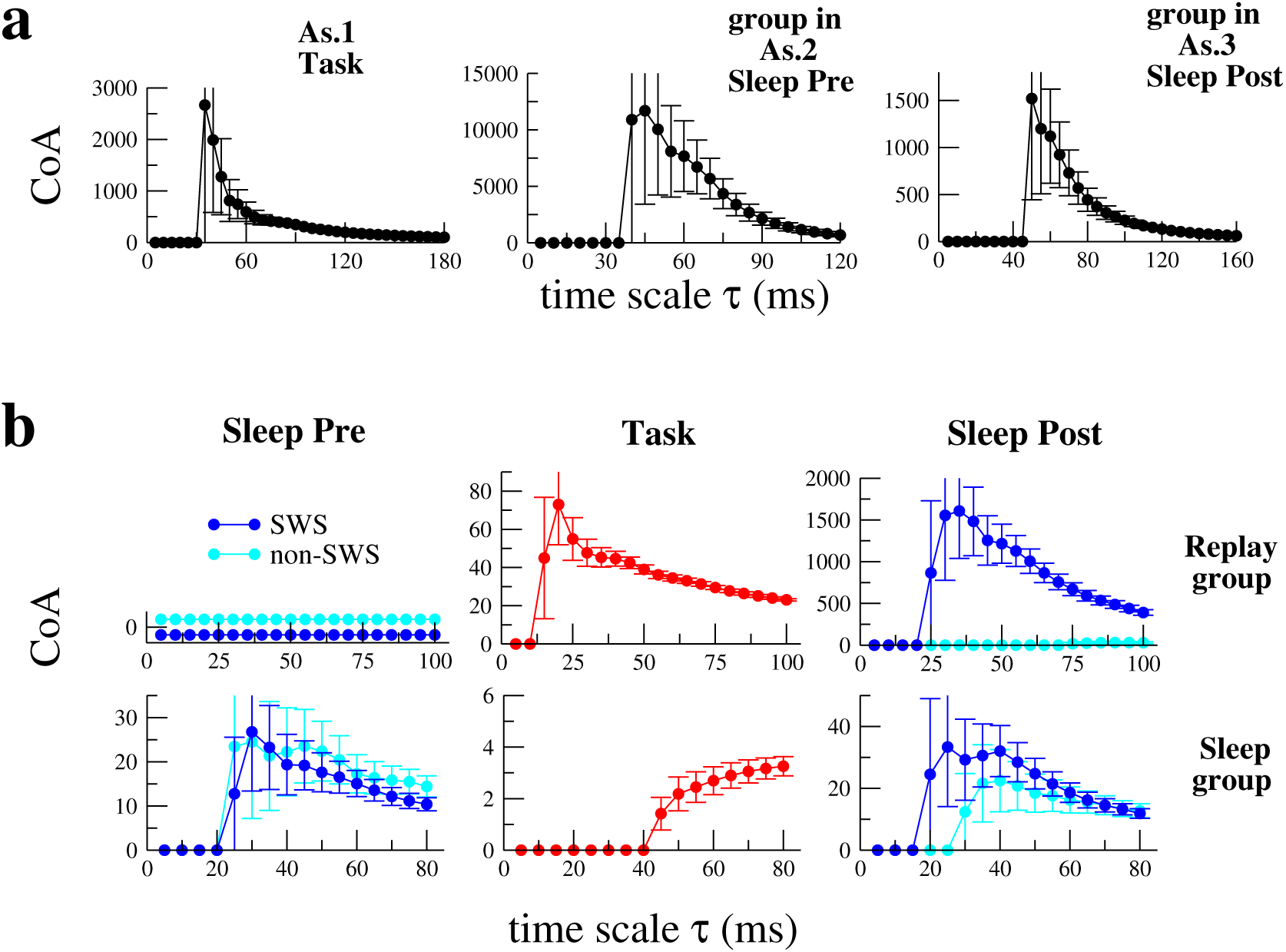
Coactivation ratios (CoA) in session A. A. CoA of As.1 in Task (*δ* log *P* ≃ 16.2), the group 2-6-9-11-12-13 in As.2 of Sleep Pre (*δ* log *P* ≃ 8.9), the group 7-8-11-12-20-21-26-35 in As.3 of Sleep Post (*δ* log *P* ≃ 12.3). **b**. CoA of the Replay and Sleep groups; see caption of Fig. 6 for the lists of corresponding neurons. CoAs are shown for time scales *τ* ranging from 5 ms to *n* × 20 ms, where *n* is the number of neurons in each group considered. Note the variations in the CoA and temporal scales along the *y*- and *x*-axis between the panels. See Methods for the computation of error bars. CoA equal to zero are compatible with the independent-cell hypothesis due to the low-firing rates of the neurons, see Methods.

The ‘Sleep’ group (neurons 6-7-11-12), shared by As.2 in Sleep Pre and the large assembly As.3 in Sleep Post, coactivates in Sleep Pre and Sleep Post, both in SWS and non-SWS periods over *τ* ≃ 30 – 40 ms (Fig. 8b). We find essentially no coactivation in Task (CoA close to 1). The ’Replay’ group (neurons 1-9-20-21-26), shared by As.1 in Task and As.3 in Sleep Post (Figs. 6a&c), coincides with the five-neuron group supporting the strongly effectively potentiated couplings identified above (Fig. 4). It strongly coactivates in Task and in SWS-Sleep Post, on similar time scales, respectively, *τ* ≃ 20 – 30 ms and *τ* ≃ 30 – 40 ms, and does not coactive in Sleep Pre nor in non-SWS periods of Sleep Post (Fig. 8b). In addition the large CoAs of the Replay group found in Task and SWS-Sleep Post are significantly higher than CoAs for random groups of five neurons (Fig. S8). Those findings support the hypothesis that the five-cell Replay group is (part of) a cell assembly involved in memory consolidation.

The coactivation of subgroups of neurons in the putative cell assemblies of each epoch can be studied further. Looking at subgroups rather than the whole assembly allows us to investigate the internal structure of the putative cell assemblies, *e.g.* the alternative activation of subgroups resulting from the presence of negative couplings. As an example, in As.2 of Sleep Pre, large CoA are found for the subgroups 2-6-9-11-12-13 (shown in Fig. 8a) and 2-6-7-11-12-13 (peak CoA value ≃ 500), but their simultaneous activation is not observed in the data. It is possible to quantitatively understand and predict which group of cells correspond to strong or to weak coactivations based on the model distribution *P*. To this purpose we introduce the log-likelihood variation (*δ* log *P*), which measures the difference in the log-likelihoods of the high and low activity configurations due to the interactions between the neurons in the group (Methods, Eqn. [5]). In session A, the only three five-cell groups found to have CoA comparable to the replay group are obtained by replacing one of the five cells in the group with another neuron in As.1; these three variants are the groups with the largest *δ* log *P* values (Supporting Information.II-A and Fig. S7). Similarly, variants of the Sleep group or subgroups of the Sleep Post assembly As.3 with large *δ* log *P* and CoAs are also identified, see Figs. 6a & S7.

### A variety of scenarios for cell assemblies are found across experimental sessions

The approach described above for session A has been applied to the other available sessions; results for six representative sessions, labelled B to G, are reported in Supporting Information-II. Given the strong and random undersampling of the neural activity it is not surprising that we find different scenarios for cell assemblies. Those scenarios are summarized and illustrated in detail on two sessions (B and D), which were recorded in two different animals, distinct from the rat of session A.

A prototypical scenario, encompassing session A, is that a Task-related group of coactivating neurons is found in Sleep Post, which was not present in Sleep Pre. An illustration is provided by session B, which consists of 10 recorded cells only. The number of active neurons in the self-sustaining configurations of the inferred Ising model of the three epochs of session B are shown in Fig. 9. No cell assembly coactivates in Sleep Pre. However, a 4-cell assembly is found in Task and is almost perfectly reproduced in Sleep Post (Fig. 9 and effective networks in Fig. 4); this cell assembly strongly coactivates in both epochs (Fig. S12). The same scenario is encountered in session G (Figs. S29 & S31).

**Figure 9:**
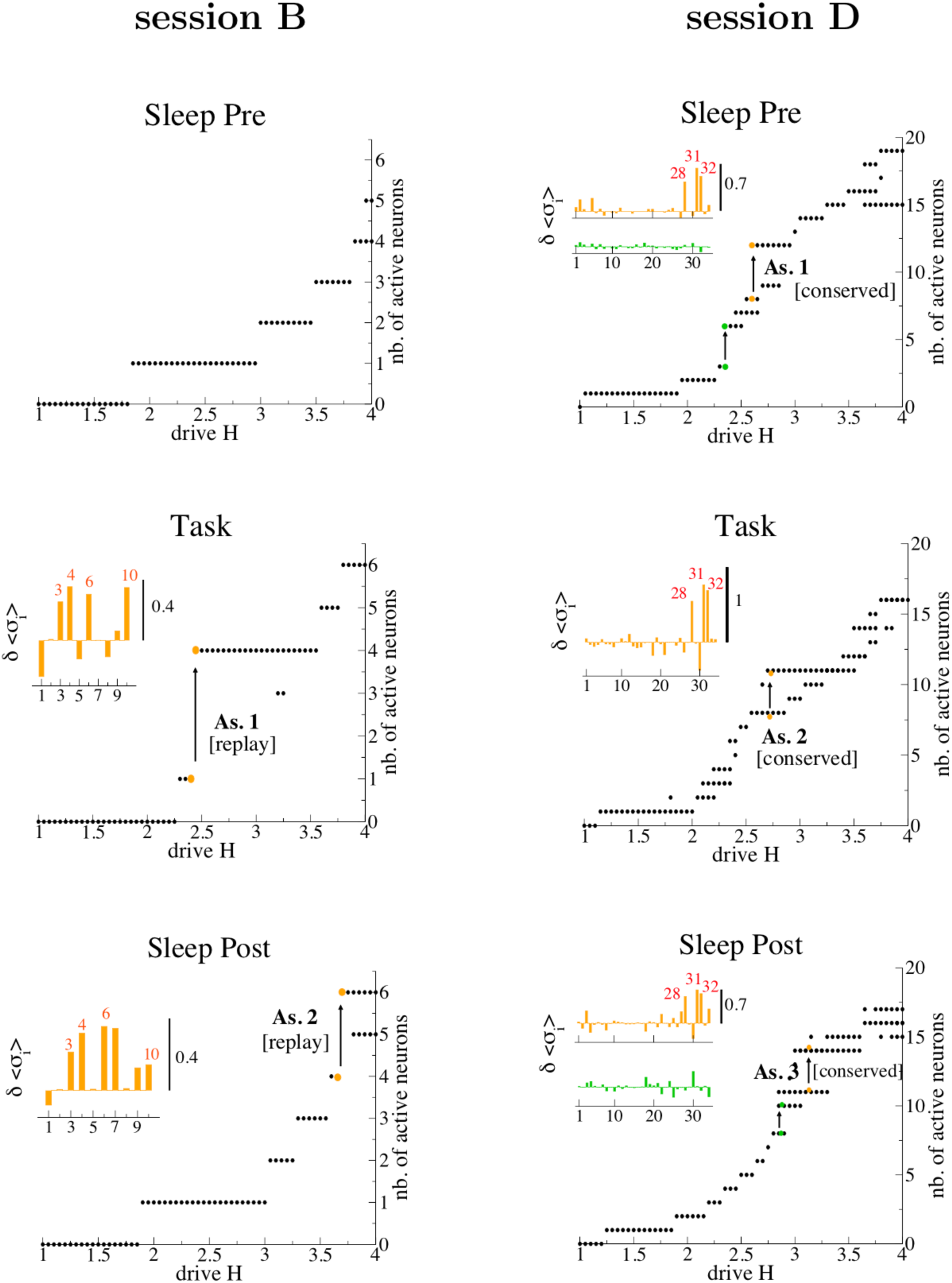
Scenarios for cell assemblies across sessions. Number of active neurons in the self-sustaining configurations of the Ising model as a function of the drive *H* for the three epochs of **sessions B** (left) and **D** (right). Coexistence of distinct self-sustaining configurations (at a given *H*) with different levels of activity (colored dots) is indicated by an arrow. The changes in the conditional averages *δ*〈*σ_i_*〉 corresponding to the jumps indicated by the arrows are shown in insets. **Session B**: Cell assemblies are found in Task and Sleep Post. Neurons of the potentiated, replay group (3-4-6-10) (indicated in orange) show a large *δ*〈*σ_i_*〉 in both Task and Sleep Post. **Session D**: Only three assemblies out of 5 jumps are found to be significant on the basis of *δ*〈*σ_i_*〉, and coincide with the group 28-31-32 (indicated in green) in every epoch. No Replay or Sleep group has been found in this session.

Other sessions are found to be even more similar to A in that they have a cell assembly in Sleep Pre, a different one in Task, and both merge in Sleep Post. An example is given by Session F, see Figs. S24, S26 & S27. In session C, the Replay group is composed of a 3-cell potentiated subgroup, whose neurons silence a 4-cell inhibited group, see Fig. 4 and caption. Remarkably, the complex structure of the Replay group is apparent through a decrease of the CoA upon addition of inhibited neurons to the potentiated 3-cell group, compare Figs. S16 & S17.

In some sessions, the same cell assembly is encountered across all three epochs. This scenario is illustrated by session D, see Fig. 9. This ‘conserved’ cell assembly is supported by three neurons (effective network in Fig. 4). We find similar values for *δ* log *P* (= 5, 6.7,6.5) and for the maximal CoA (= 25, 40,40) in, respectively, Task, Sleep Pre and Sleep Post, see Fig. S20.

Many sessions show very small values of the Adjustment *Adj*, Eqn. [3] and Fig. 5, and show no coactivation at all or a conserved cell assembly as in session D above. Interestingly, a few sessions with large or intermediate values of *Adj* do not exhibit any cell assembly; for those sessions, the effectively potentiated couplings do not interconnect a small set of recorded neurons, but are scattered over non-overlapping pairs of neurons. An example is provided by session E, see Fig. S22.

## Discussion

As a conclusion, our Ising-model approach offers a natural way to detect and study cell assemblies in terms of the (co)activation properties of a virtual neural network. Those assemblies can be related to task learning or not, as the ones which are specific to the Sleep epochs. In the former case, our method can detect replay events, even when ‘templates’ [7, 8, 9] as provided for example by the sequential activation of hippocampal place cells, are not available. Our analysis significantly extends the principal component analysis (PCA) of [10, 11], as it identifies the neurons participating to cell assemblies in a detailed way in all epochs. The largest entries of the top components of the Pearson correlation matrix have some correspondence with the neurons in the coactivated groups identified with the present method, especially in the Task epoch. However, disentangling those co-activated groups is difficult, even with the use of clustering procedures [12], see Supporting Information III for more details.

The approach consists in three steps: (1) The inference of couplings for each epoch allows us to compute the coupling Adjustment *Adj* for each session, and retain large-*Adj* sessions for the study of experience-related replay; (2) The simulation of the inferred models under an external drive *H* permits us to determine coactivating groups of neurons, and to assess their robustness through their log-likelihood variation *δ* log *P*; (3) The coactivation properties of these putative cell assemblies are then directly estimated from the spiking data through their CoA. Steps 1&2 complement and validate each other. Large coupling Adjustments can be found without experience-related cell assembly and replay, *e.g.* when potentiated couplings do not sufficiently interconnect the recorded neurons, as in session E. Similarly, cell assemblies mixing coactivation and inhibition (as in session C) would be hard to characterize from the ’jump’ analysis only.

During wakeful experience and during sleep, neural activity shows very different regimes. The Ising models can bridge the gap between these states by clearly separating coupling and input effects. As the firing rates are, on average, very similar in Sleep Pre and Sleep Post, the creation of task-related cell assemblies mainly result from a change in the effective couplings *J_ij_*, which we quantify with the coupling Adjustment *Adj*, Eq. [3]. Whether *Adj* is related to physiological, synaptic plasticity or not is a fascinating and totally open question. A difficulty, intrinsic to multi-electrode recordings, is that the random and limited sampling may miss important cells, and skew our estimate of experience-related changes to the couplings. This may partially explain the presence of session-to-session fluctuations in *Adj* (Fig. 5).

A crucial ingredient of our approach is the addition of a global drive *H* to the local inputs *h_i_* in order to detect the cell assemblies (Fig. 6). The set of *h_i_* estimated from the data allow us to fit the diversity of session-averaged firing rates for different neurons, but cannot reproduce transient fluctuations in activity, which are important for cell assembly recruitment. It is a remarkable fact that ’rare’ coactivation events can be evoked through an extra-stimulation of the model inferred from the session-average activity. It would be interesting to study whether this result holds with other inference approaches, *e.g.* generalized linear models [16] or integrate-and-fire models [18, 19]. In physiological terms, *H* may translate to increased neuronal excitability, synaptic facilitation/depression state, or large transient inputs, such as those arising from hippocampal sharp waves [10]. The choice of a homogeneous input *H* to identify the groups of coactivating neurons ensures that the coactivating neural groups found are induced by the parameters fit on the data (set of *J_ij_* and *h_i_*) only. In addition, non-homogeneous inputs targeting one or more neurons eventually reveal the same neural groups in case of strong coactivation (Supporting Information-II.A, Fig. S9).

Our approach to identify cell assemblies is based on the notion of simultaneous coactivation [4], irrespectively of temporal ordering aspects. The network of pairwise couplings inferred on short time scales, Δ*t* = 10 ms, suffices to predict coactivation patterns between *n* neurons on longer time scales, *τ* ≃ *n* × Δ*t* ms. Informally speaking the repeated spiking of neurons in short ‘bursts’ allows for the coverage of *τ* with a sequence of pairwise coactivation events (Fig. 1). As the Ising model gives the distribution of snapshots of the activity couplings are symmetric, and no ordering can be predicted by the model. Yet, we have not been able to identify a clear activation ordering in the replay groups in the spiking data. A possible explanation is that the information encoded in prefrontal cortex has, indeed, aspects that are not inherently sequential, in particular, the current rule used to solve the task.

Simple empirical rules for cell assembly modification emerge from the analysis of the various experimental sessions. If a cell assembly is found in Task, while no coactivation is seen in Sleep Pre, then this cell assembly is found also in Sleep Post, *e.g.* Session B in Fig. 9. If cell assemblies are found both in Task and in Sleep Pre, but those assemblies differ in their constituting neurons, then they become associated in Sleep Post, e.g. the merging of jumps in session A (Figs. 6 & 7). This association can be accompanied by a reshaping of cell assemblies in the presence of effective inhibitory couplings, e.g. in session C. Last of all, if the same cell assembly is encountered in Task and in Sleep Pre, then it is conserved in Sleep Post, see session D in Fig. 9. In regards to this empirical rules, the large body of knowledge on Ising models with non-homogeneous couplings accumulated over the last decades [30] could prove useful to improve our theoretical understanding of how cell assemblies can be created, modified, suppressed, or combined with each other [3]. Combined with optogenetics techniques [31] this would open exciting perspectives in the manipulation of cortical cell assemblies in a controlled way.

## Materials and Methods

### Inference and validation of Ising distribution of activity

We associate to each neuron *i* in time bin *t* a variable *σ_i_*(*t*) = 1 if the neuron is active, 0 if it is silent. The frequencies *f_i_* and *f_ij_* are defined as the average values over time bins of, respectively, the variables *σ_i_*(*t*) and *σ_i_*(*t*)*σ_j_*(*t*). We look for the Ising model parameters {*h_i_*, *J_ij_*} such that *f_i_* and *f_ij_* match, respectively, the average values of *σ_i_* (for all neurons *i*) and *σ_i_σ_j_* (for all pairs of neurons *i, j*) over *P*, Eqn. [1]. The inference of the parameters is carried out using the Adaptive Cluster Expansion (ACE) of [26, 29], see Supporting Information-I.A. ACE recursively builds up of clusters of strongly interacting neurons of increasing sizes, whose contributions to the entropy of the Ising model are larger than a threshold Θ. The value of Θ is chosen to reproduce accurately the single-neuron (*f_i_*) and pairwise (*f_ij_*) spiking probabilities, see Fig. 2a and Supporting Information-I.A, within the expected uncertainty due to the finite recording time. Statistical errors bars {Δ*h_i_*, Δ*J_ij_*} on the inferred parameters are also computed, see Supporting Information-I.B.

To compute the average values of observables with the inferred model distribution *P*, we resort to Monte Carlo simulations. The quality of the reproduction of the single-neuron and pairwise spiking probabilities in a time bin is shown in Fig. 2a, and the predictions of higher-order moments, such as triplet firing probabilities and the probability of multiple-neuron firing in a time bin, are compared to the data in Fig. 2b. To cross-validate the model *P* we divide the data set of the Task epoch of the session A in two halves. We extract the spiking probabilities from the first half of the recording, and we infer the Ising model able to reproduce these data within their statistical errors. We then compare in Fig. 2c the correlations *c_ij_ = f_ij_ – f_i_f_j_* obtained with the model (through Monte Carlo Markov Chain simulations) to their experimental counterparts computed from the first half (used for the inference, left panel), and from the second half (independent of the inference, right panel) of the data. We choose to represent the correlations *c_ij_* rather than the pairwise probabilities *f_ij_* as the former are more sensitive to errors in the inference than the latter. The excellent agreement confirms the absence of overfitting in our inference.

### Statistical significance of the Coactivation factor (CoA)

To assess the statistical validity of the CoA defined in Eq. [4] for a group *G* of neurons we compute the error bar on CoA, shown in Fig. 5. Assuming a Poisson distribution for the co-activation events, the standard deviation of the CoA is estimated to be 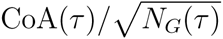, where *N_G_*(*τ*) is the number of coactivation events for the cells in *G* over the time scale *τ*. Note that simultaneous-firing events (contributing to *f*(*G*)) are unlikely to be found, and the CoA is likely to be zero, if the duration of the recording is small, *e.g.* compared to 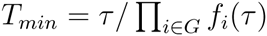. This happens for the five-cell replay group of session A for time scales *τ* ≤ 55 ms in the Task epoch, and for all the values of *τ* considered in Sleep Pre and Post in Fig. 8.

### Search for self-sustaining neural activity configurations

We start by showing that self-sustaining configurations ***σ*** = (*σ*_1_ *σ*_2_,*…, σ_N_*) are in one-to-one correspondence with the local maxima of the distribution P, Eq. [1]. Consider a neuron, say, *i*, and its total input *V_i_*, given by Eq. [2]. By definition of a self-sustaining configuration, we have either *σ_i_* = 0 and *V_i_* < 0, or *σ_i_* = 1 and *V_i_* > 0. In the latter case, after changing the activity of neuron *i* from *σ_i_* = 1 to *σ_i_* = 0 the log-probability log *P* decreases by *V_i_* > 0. In the former case, after changing the activity of neuron *i* from *σ_i_* = 0 to *σ_i_* = 1 the log-probability log *P* increases by *V_i_* < 0. In both cases we see that changing the value of the neuron activity leads to a decrease of log *P*.

The search procedure for configurations in which all neurons are self-sustaining is the following. We start with the all-silent neuron configuration (*σ_i_* = 0 for *i* = 1,…, *N*). If the configuration is self-sustaining, the algorithm has found a maximum of *P* and halts. If one or more neurons are not self-sustaining, *i.e.* their values *σ_i_* do not agree with the signs of their total inputs *V_i_*, we pick up uniformly at random one of them, say, *i*, and flip its value *σ_i_* (from silent to active, or vice-versa). This asynchronous updating is iterated until the configuration is self-sustaining. The procedure is guaranteed to converge as the log-probability of the configuration increases after each updating step. Re-running the dynamics may, however, produce different maxima, due to the stochasticity in the choice of the (non self-sustaining) neuron to flip at each step.

### Cell assemblies, and log-likelihood variation

For each value of the drive *H* we determine the self-sustaining configurations ***σ***, their numbers of active neurons, and the total inputs *V_i_*(***σ***) + *H* to those active neurons, see paragraph above. A jump corresponds to the coexistence of two self-sustaining configurations ***σ***^(1)^ and ***σ***^(2)^ for the same value of *H*, with more neurons active in ***σ***^(2)^ (high activity) than in ***σ***^(1)^ (low activity). To determine the cell assembly attached to the jump we rank all the cells *i* according to the variation 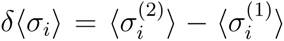 of their conditional average values, see Eq. [2]. Neurons with the largest *δ*〈*σ_i_*〉 are included in the assembly. The cut-off value over *δ*〈*σ_i_*〉 is chosen to be ≃ 0.2, but may depend on the session, see Supporting Information-II.

We define the *log-likelihood variation* of the cell assembly as the change in log-likelihood due to the interactions only:

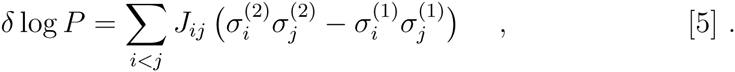

The log-likelihood variation of a subgroup of neurons in ***σ***^(2)^ is defined as in Eq. [5], with the double sum restricted to the neurons in the subgroup.

## Supporting Information Files

**Text S1**. Supporting Information for Inferred Model of the Prefrontal Cortex Activity Unveils Cell Assemblies and Memory Replay.

## Acknowledgments

This work was funded by the [EU-]FP7 FET OPEN project Enlightenment 284801.

